# Pichinde virus models intrauterine infection by hemorrhagic fever-causing arenaviruses

**DOI:** 10.64898/2026.09.15.751927

**Authors:** Craig J. Bierle, Hannah Murphy, Jason S. Hatfield, Qinfeng Huang, Tyler B. Rollman, Priyanka Chauhan, Brigitte Flannery, Sarah A. Wernimont, Terry K. Morgan, Micah D. Gearhart, Yuying Liang, Hinh Ly

## Abstract

Arenavirus infection during pregnancy can cause severe maternal disease, congenital infection, and fetal demise. Many arenaviruses are endemic in economically disadvantaged areas and must be studied in high containment laboratories, which has limited our understanding of arenavirus pathogenesis in the placenta. Pichinde virus (PICV) is a nonpathogenic arenavirus that recapitulates many aspects of viral hemorrhagic fever in guinea pigs. Using a combination of experiments in human placental cells and guinea pigs, we characterized the tropism and impact of PICV infection during pregnancy. PICV replicated in human trophoblast stem cells (TSCs), TSC-derived trophoblasts, and explanted term placenta. When guinea pigs were infected at mid-gestation, PICV caused fetal demise. High infectious titers were recovered from placenta and decidua, but PICV was infrequently detected in fetal tissues or amniotic fluid. *In situ* hybridization confirmed that the placenta, decidua, and fetal membranes were all infected by the virus. Transcriptional profiling of PICV-infected human trophoblasts and guinea pig tissues revealed that infection upregulated canonical antiviral responses, which could contribute to placental dysfunction and pregnancy loss. Thus, PICV is safe and tractable model of zoonotic arenavirus infection during pregnancy with utility for preclinical therapeutic and vaccine development.

**Author Summary:** The arenaviruses are emerging pathogens that can cause high mortality viral hemorrhagic fevers. Several of these zoonotic viruses cause severe infections in pregnant people that are associated with high rates of maternal death, pregnancy loss, and disability in affected children. The placenta is normally an effective barrier against the mother-to-child transmission of bloodborne viruses, but arenavirus replication in the placenta may contribute to severe maternal disease and adverse fetal outcomes. We demonstrate that Pichinde virus, which is not a human pathogen, can replicate in human placental cells and cause pregnancy loss in guinea pigs, establishing a new experimental model for understanding viral hemorrhagic fever in pregnancy.

## Introduction

The arenaviruses are a family of emerging viruses and potential pandemic pathogens. Zoonotic transmission of arenaviruses from their rodent hosts to humans can cause high-mortality viral hemorrhagic fever (VHF) [1]. Arenaviruses and other pathogens that cause VHF are significant but under-recognized causes of maternal and fetal morbidity and mortality in Africa, Central, and South America, and Asia [2]. Three arenaviruses have been noted to cause significant morbidity and mortality in pregnant humans or in animal models. Lassa virus (LASV) infection causes maternal mortality rates up to 50% and pregnancy loss in 75-92% of cases [2-5]. LASV replication at the maternal-fetal interface may cause these adverse outcomes as high viral loads are detected in infected placenta and maternal outcomes can rapidly improve when the uterus is evacuated [4, 6]. Lymphocytic choriomeningitis virus (LCMV) is less virulent than LASV but can cause debilitating congenital infections. Brain abnormalities, such as chorioretinitis, hydrocephaly, and microcephaly, occur in 97% of congenitally-infected fetuses, and intrauterine infection often results in fetal demise [7]. Junin virus (JUNV), the cause of Argentine hemorrhagic fever (AHF), can cause high rates of mortality, abortion, and placental infection in guinea pigs [8, 9]. A case series of twelve women diagnosed with AHF during pregnancy reported high rates of maternal mortality after infection in the third trimester [10]. The vertical transmission of arenaviruses *in utero* or early in postnatal life from persistently infected dams to their offspring plays a critical role in maintaining the pathogens in their rodent reservoirs [1]. Despite the importance of placental infection in the natural history of arenaviruses and in disease following zoonotic infection, little is known about the pathogenesis of arenaviruses during pregnancy and how viral replication at the maternal-fetal interface affects clinical outcomes. LCMV establishes persistent infections in mouse placenta and replicates to high titers in explanted first-trimester human placenta but poorly in term tissues [11-13]. LASV antigens are detected in infected placenta, but the virus can only be studied in high-containment BSL-4 facilities, and experimental infections of placental cells, tissues, or pregnant animals have not been reported [14].

Pichinde virus (PICV), an arenavirus that was originally isolated from Tomes’s rice rat (*Nephelomys albigularis*), is non-pathogenic in humans and has been used as a low-biosafety surrogate model of VHF in guinea pigs [15-17]. To establish this model, PICV was serially passaged in inbred strain 13 guinea pigs. Low passage (P2) virus is avirulent whereas high passage (P18) virus is highly pathogenic and causes VHF-like illness in the rodent [18, 19]. Guinea pigs are a powerful model for understanding the developmental origins of health and disease, with placental physiology and intrauterine development that is more representative of primates than murid rodents [20, 21]. To establish a model of zoonotic arenavirus infection during pregnancy, we evaluated whether PICV could infect trophoblasts and cause adverse pregnancy outcomes in guinea pigs. We found that PICV replicated in explanted term human placenta, where the virus was most frequently detected in syncytiotrophoblasts (STBs), and in human trophoblast stem cells (TSCs) and TSC-derived extravillous trophoblasts (EVTs) and STBs. When pregnant guinea pigs were infected at mid-gestation, both P2 and P18 caused fetal demise, high PICV titers in the placenta and decidua, and infrequent fetal infections. Transcriptional profiling of PICV-infected TSCs and guinea pig tissues revealed gene expression changes indicative of antiviral signaling, including type I interferon responses, that may contribute to placental dysfunction [22, 23]. Thus, like LASV, LCMV, and JUNV, PICV has tropism for the placenta and may be a valuable preclinical model for understanding how VHF causes adverse pregnancy outcomes.

## Results

### PICV replicates in term human placenta, trophoblast stem cells, and differentiated trophoblasts *in vitro*

The placenta and fetal-derived trophoblasts are broadly antiviral and refractory to many viral infections [24, 25]. To assess whether PICV could infect the human placenta *in vitro*, chorionic villi explants were prepared from tissue collected at term and infected with PICV P18 (5x10^5^ PFU/explant). Cell culture media and tissue explants were collected every 24 h and viral loads in the media and tissue homogenates were assessed by plaque assay (**Fig. 1A**). Infectious PICV was recovered from both sample types, and the abundance of virus increased 10- to 100-fold between 24 and 72 hours post infection (hpi). *In situ* hybridization targeting the PICV nucleoprotein (*NP*) gene, which detects both viral mRNA and antigenomes, was used to visualize infected cells in the tissue explants (**Fig. 1B**). At 24 hpi, *NP* was almost exclusively detected in STBs. PICV spread extensively through the STB layer between 48 and 72 hpi, and individual *NP* transcripts could be detected in the core of villi at these later times.

**Figure 1.**
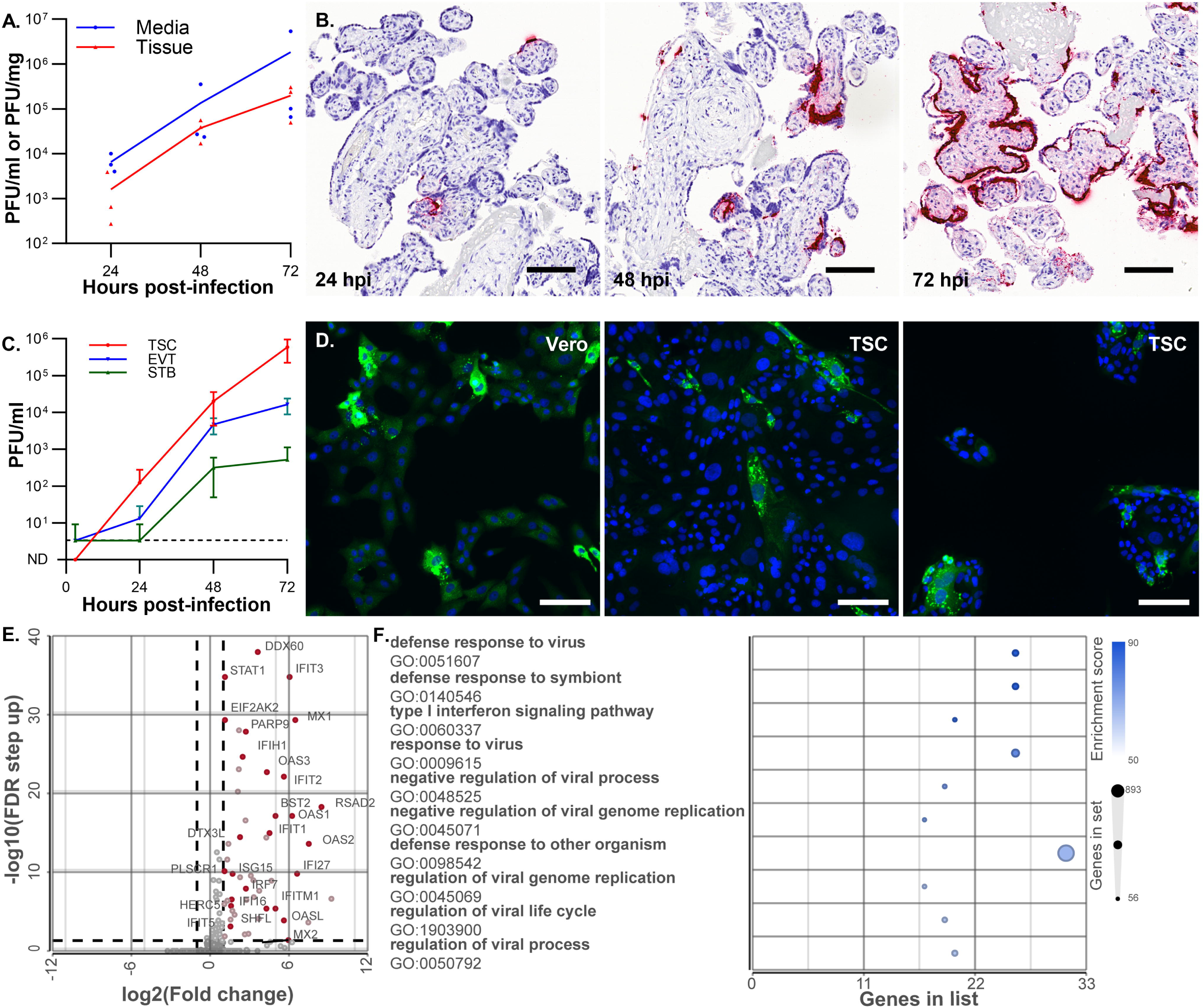
PICV replicates in human trophoblasts. Chorionic villi explants were prepared from term human placentas and infected with PICV. (**A**) Viral loads in media and explants were determined by plaque assay every 24 h. Data from three experiments, each completed using a unique placenta, is shown, and each point represents the mean titer of three explants. (**B**) PICV-infected explants were formalin-fixed and paraffin-embedded. *NP*-specific RNAscope was used to detect infected cells (scale = 100 µm). (**C**) TSCs were differentiated into EVT and STB, and the three cell types were infected with PICV (MOI=0.1). Infected cells were scraped into their media, and the abundance of virus was determined by plaque assay. (**D**) Vero cells and TSCs were infected with PICV (MOI=10), fixed, and stained with α-PICV polyclonal antibody and DAPI at 24 hpi (scale = 100 µm) (**E**) TSCs were infected with PICV (MOI=10) or mock-infected. RNA was extracted and cellular transcript abundance was assessed by RNA-Seq. Transcripts that were differentially regulated by infection are illustrated by volcano plot (51 upregulated, ≥2-fold, FDR ≤ 0.05); transcripts related to defense response to virus (GO:0051607) are emphasized. **(F)** The ten GO terms that were most highly enriched among transcripts regulated by PICV infection, as determined by gene set enrichment analysis, are illustrated by bubble plot.

TSCs were used as a second *in vitro* model to confirm that PICV can infect and replicate in human trophoblasts. These cells are phenotypically representative of first trimester trophoblasts and can be maintained in an undifferentiated state or differentiated into EVT or STB [26-28]. Undifferentiated TSCs and TSC-derived EVT and STB were infected with PICV P18 at a multiplicity of infection (MOI) of 0.1, and the abundance of PICV was measured by plaque assay at 3, 24, 48, and 72 hpi (**Fig. 1C**). PICV replicated in all three cell types, but more virus was recovered from the undifferentiated TSCs than the differentiated EVTs or STBs. To visualize infection in TSCs, Vero cells and TSCs were infected with PICV P18 (MOI=10) and stained with PICV-specific polyclonal antibodies at 24 hpi (**Fig. 1D**). While both cell types expressed viral antigens, the proportion of antigen-positive cells was lower than expected in a high multiplicity infection. To determine how PICV infection affects gene expression in TSCs, RNA was extracted from mock- and PICV P18-infected cells (MOI=10) at 24 hpi and sequenced. Fifty-one cellular transcripts were significantly upregulated by PICV infection (≥2-fold, FDR < 0.05) (**Fig. 1E**). Gene set enrichment analysis found that many pathways related to antiviral defense, including defense response to virus (GO:0051607), the type I interferon signaling pathway (GO:0060337), and regulation of viral life cycle (GO:1903900) were highly enriched in the infected cells (**Fig. 1F**). While interferon expression was not significantly modulated by the infection, transcripts encoding many interferon-stimulated genes (ISGs) were upregulated by the infection.

### PICV causes fetal demise in guinea pigs but infrequently transmits to the fetus

Guinea pigs have been frequently used to study the pathogenesis of VHF. Clinical LASV isolates and guinea pig-adapted PICV P18 can cause fatal infections in inbred strain 13 guinea pigs [18, 19, 29-31]. Guinea pigs have 65-70 day gestational periods, and the intrauterine development of their major organs and central nervous system is highly similar to primates [21]. Guinea pigs have primate-like hemomonochorial placentas, wherein a single layer of syncytiotrophoblasts separates maternal blood from fetal circulation, and have been used to model intrauterine infections caused by diverse human pathogens [20, 32-37]. To assess how PICV infection affects pregnant guinea pigs, time-mated strain 13 dams were infected with low (P2) and high (P18) virulence PICV variants [17].

Groups of time-mated and non-pregnant females were infected with P18, and three groups of pregnant animals were infected with PICV P2 (N=3/group). The pregnant guinea pigs were infected at mid-gestation (35 days gestation [dGA]); this was during the fetal period when dams rapidly gain weight under normal conditions (**Fig. 2A, S1 Fig.**) [38-40]. Infected non-pregnant animals met our endpoint criterion of 15% weight loss at 10 dpi and were euthanized. All dams infected with P18 lost weight relative to day zero (D0) and delivered stillborn pups at either 9 or 10 dpi. When compared to dams that had been mock-infected with saline at 35 dGA in previous studies, P2-infected dams did not gain the expected amount of weight after infection [39]. No P2-infected dam delivered before their scheduled experimental endpoint and groups of animals were euthanized at 10, 15, and 20 dpi. The sizes of fetuses and placentas from PICV-infected dams were compared with historical data from mock-infected guinea pig pregnancies that ended at 42, 49, and 56 dGA (**Fig. 2B, C and Table S1**). Fetal guinea pigs normally grow between 0.6 and 2.7 g per day after mid-gestation. Differences in guinea pig strain, gestational age, and litter size, which all could affect the size of fetuses and their placentas, confound our ability to make comparisons between PICV infected pregnancies and our historical data [20, 41]. Nonetheless, stillborn pups recovered after P18 infection at 9-10 dpi (44-45 dGA) were significantly smaller than normal 42 dGA fetuses and age-matched fetuses from P2-infected dams. The size of fetuses and their placentas did not appear to be affected by P2 infection at either 10 or 15 dpi. At 20 dpi, 2 of 3 P2-infected litters had succumbed to infection *in utero*; these fetuses were undersized and, upon evaluation of their body condition, resorbing. The third litter in this group appeared normal, although one placenta was noted to be hemorrhaged upon gross examination.

**Figure 2.**
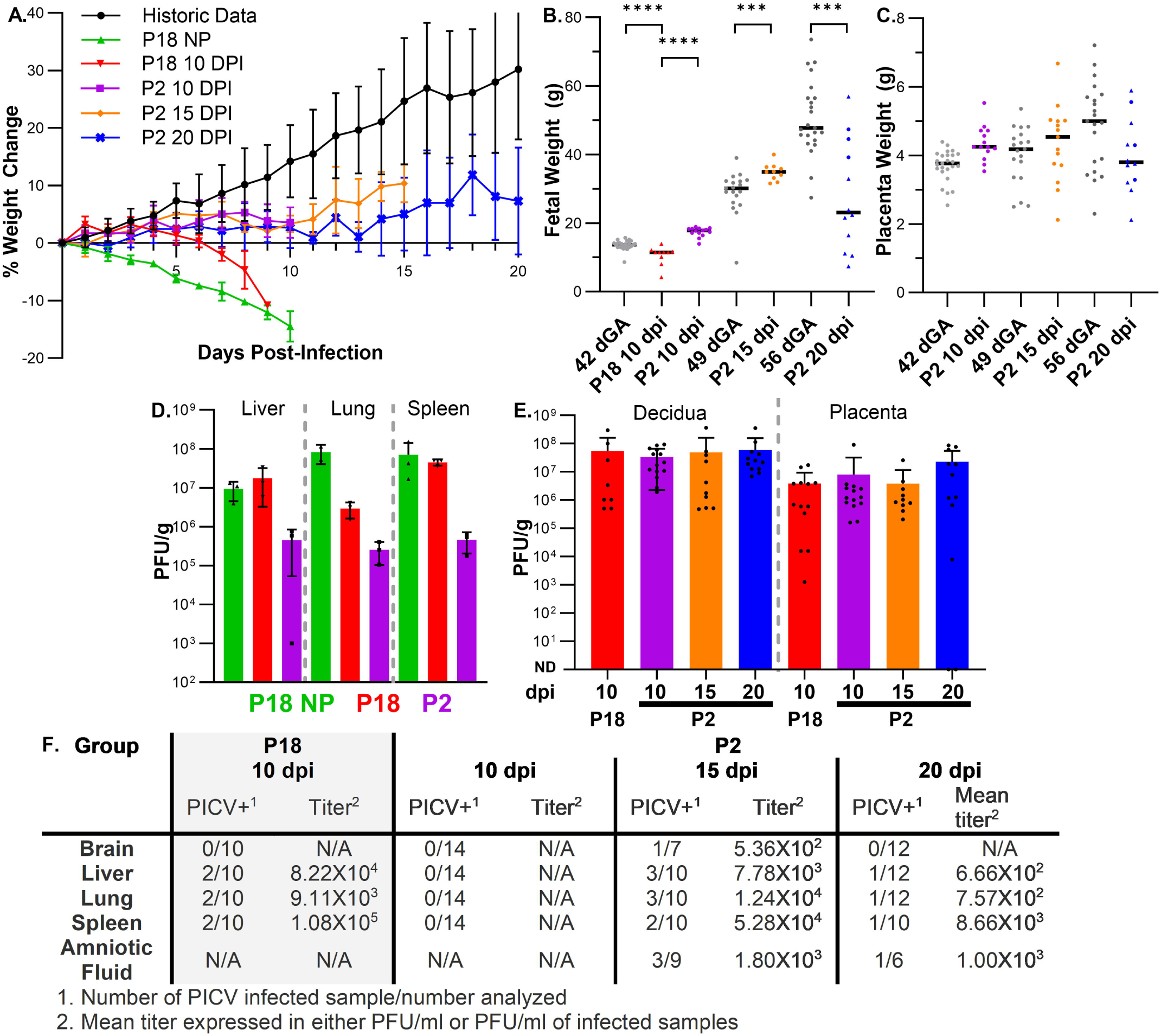
PICV affects maternal health and infects the maternal-fetal interface in guinea pigs. Time-mated and non-pregnant (NP) guinea pigs were infected with PICV P2 or P18. (**A**) Weight change post-infection was plotted alongside historical data, which illustrates normal weight gain in pregnant guinea pigs between 35 and 55 dGA. Fetal weight (**B**) and placental weight (**C**) after stillbirth or at necropsy plotted with reference data from mock-infected samples collected at 42, 49, and 56 dGA. Triangles indicate pups that were stillborn or found dead at necropsy (*** *p* < 0.001, **** *p* < 0.0001, Mann– Whitney U test). Plaque assay was used to determine PICV viral loads in the maternal lung, liver, and spleen 10 dpi (**D**), in decidua and placenta (**E**), and in fetal tissues and amniotic fluid (**F**).

PICV viral loads in tissue homogenates and amniotic fluid were quantified by plaque assay. At 9-10 dpi, P2 and P18 PICV were both detected in maternal viscera; viral loads were higher in the P18-infected samples (**Fig. 2D**). Virus was not detected in the maternal lung, liver, or spleen of the P2-infected dams at 15 or 20 dpi. Both P2 and P18 caused high viral loads in the decidua and placenta (**Fig. 2E**). However, the virus was only occasionally detected in fetal remains or amniotic fluid (**Fig. 2F**). Virus was typically recovered from multiple tissues from PICV-infected fetuses, but viral loads in fetal tissues were markedly lower than what was observed in the placenta or decidua. Taken together, these results reveal that PICV infects the maternal-fetal interface but only occasionally transmits to the developing fetus.

### PICV infection causes placental inflammation, necrosis, and infarction

To understand how PICV infection affects the guinea pig placenta, infected placentas were sectioned, hematoxylin and eosin (H&E) stained, and analyzed by a perinatal pathologist. Findings are summarized in **Table S1**. After P18 infection, the majority of placentas (9 of 13) contained a pattern of acute inflammation consistent with infection. Diagnostic abnormalities were uncommon in P2-infected placentas collected at either 10 or 15 dpi. Acute placental infarction was observed in 1 of 14 samples examined at 10 dpi; 2 of 10 placentas examined at 15 dpi had evidence of necrosis in the labyrinth. In contrast, the majority of P2-infected placentas (9 of 12) had one or more abnormal pathologic findings at 20 dpi. These findings include infarction, necrosis and/or calcification of the labyrinth (7 of 12), and death of the parietal yolk sac epithelium (4 of 12).

Two placentas from each litter were further analyzed by PICV-specific RNAscope to visualize the location viral RNA in the tissue and the extent of infection. Viral RNA was detected at the maternal-fetal interface in all P18-infected samples, but the pattern and intensity of staining was variable (**Fig. 3**). P18 PICV transcripts were detected in the decidua and parietal yolk sac in all samples analyzed (6 of 6, **Fig. 3B**). P18 transcripts were occasionally detected in the subplacenta. In several samples, individual viral transcripts or infected cells were seen throughout the labyrinth. In two of the samples large areas of viral *NP* staining were observed in necrotic regions of the labyrinth (**Fig. 3C**). In comparison to P18, P2 PICV appears to have progressed through the placenta with slower kinetics. At 10 dpi, intense *NP* staining was observed in the decidua and parietal yolk sac (**Fig. 3D-F**), and P2 viral RNA was occasionally detected in the subplacenta and/or labyrinth. High amounts of P2 viral RNA were observed in the parietal yolk sac at 15 dpi, but *NP* abundance was markedly diminished in the decidua and the transcript was rarely seen in the syncytium (**Fig. 3G-I**). The fewest P2 transcripts were detected in the decidua at 20 dpi (**Fig 3. J-L**), but viral transcripts remained abundant in parietal yolk sac, and most samples (4 of 6) contained infected cells throughout the labyrinth.

**Figure 3.**
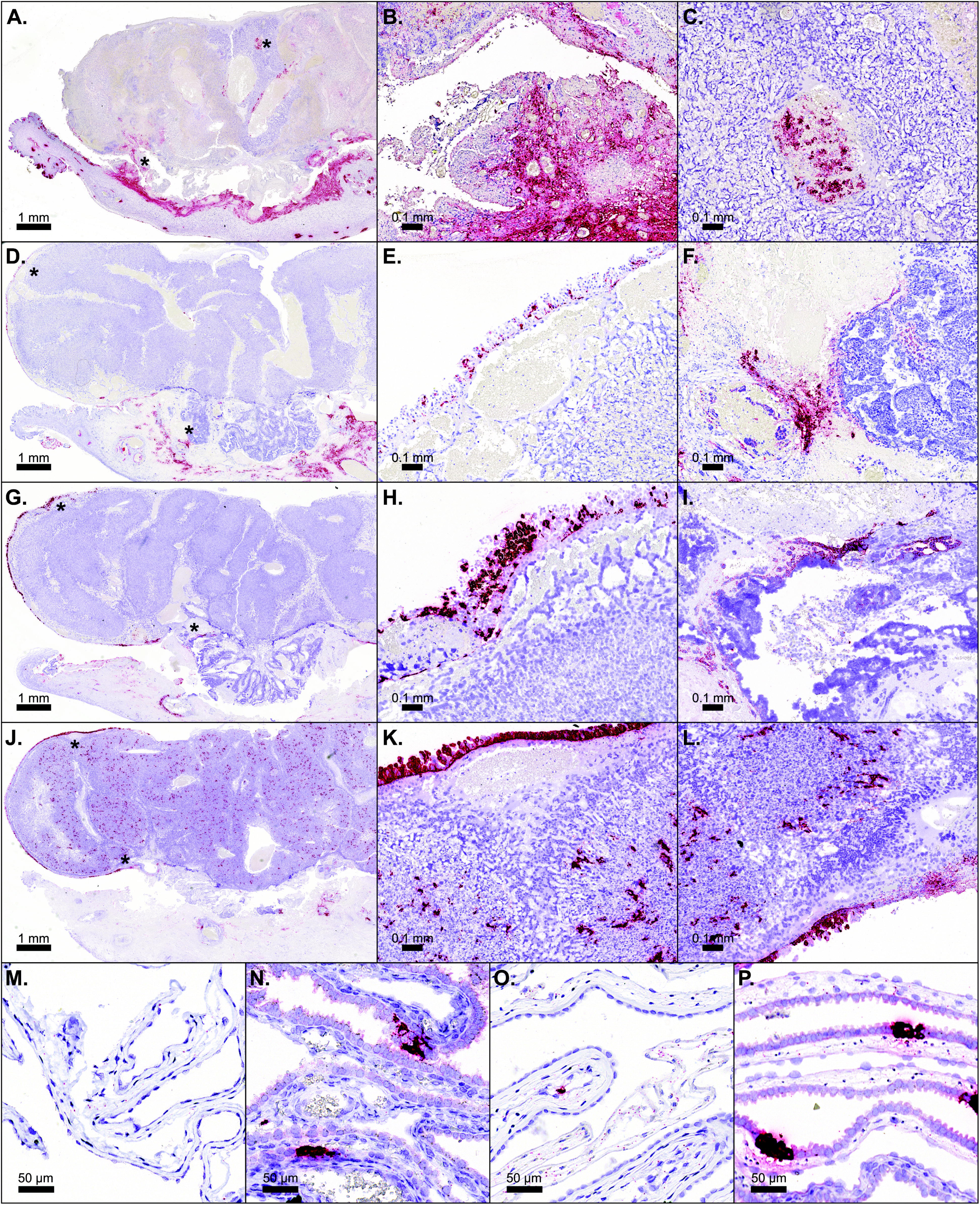
PICV RNA is detected at the maternal-fetal interface. Formalin-fixed, paraffin-embedded placentas and fetal membranes were stained by *NP*-specific RNAscope. Representative samples are shown; asterisks indicate high magnification regions of interest from each sample. P18-infected placenta at 10 dpi (**A**) highlighting PICV-infected cells in the junctional zone (**B**) and labyrinth (**C**). P2-infected placenta at 10 dpi (**D**); high magnification images show infected cells in the parietal yolk sac epithelium (**E**) and in the junctional zone and subplacenta (**F**). P2-infected placenta at 15 dpi (**G**) with representative images illustrating PICV infection in the yolk sac epithelium (**H**) and subplacenta (**I**). P2-infected placenta at 20 dpi (**J**) with high magnification images of infected cells in the yolk sac epithelium and labyrinth (**K, L**). PICV P2-infected amnion (**M**) and visceral yolk sac (**N**) at 15 dpi. P2-infected amnion (**O**) and visceral yolk sac (**P**) at 20 dpi. Full resolution slide scans for all samples analyzed in this study are available for download at BioImage Archive under ascension S-BIAD3362.

Having noted that PICV infected the parietal yolk sac, which surrounds the placenta, samples of the amnion and the visceral yolk sac were collected from each animal in the P2-infected 15 and 20 dpi groups and analyzed by H&E staining and RNAscope (**Fig. 3M-P**). *NP* staining was seen in all samples of the visceral yolk sac examined at 15 dpi. One of three samples of amnion contained *NP*^+^ fibroblasts in the subamniotic mesoderm. Neither membrane had other diagnostic abnormalities at the time point. However, all three samples of visceral yolk sac that were examined were necrotic at 20 dpi and contained PICV-infected epithelial cells and macrophages. Viral transcripts were detected throughout all the samples of amnion examined, but few amnion cells were intensely stained with the *NP* probe.

### Transcriptional profiling of the PICV-infected maternal-fetal interface

To illuminate how PICV infection affects the guinea pig maternal-fetal interface, RNA was extracted from placenta and decidua and sequenced. For this analysis, samples from P2- and P18-infected animals were collected at 9 or 10 dpi. Gestation age-matched healthy tissue was included as a normal control. Comparing normal and P18-infected samples, more transcripts were differentially regulated (≥ 2-fold, FDR ≤ 0.05) by infection in the placenta than in the decidua (**Fig. 4A**). Gene set enrichment analysis found that transcripts related to antiviral immunity, including the terms immune system process (GO:0002376) and defense response to virus (GO:0051607), were enriched and generally upregulated by infection in both tissue compartments. Other notable findings in P18-infected placenta include that genes related to lipid metabolic process (GO:0006629) were downregulated and transcripts related to a response to wounding (GO:0009611) were upregulated. P2 infection had comparably little effect on host gene expression at 10 dpi, and few transcripts (3 in decidua and 18 in placenta) met the FDR ≤ 0.05 cutoff for differential regulation when infected and normal tissue was compared. When transcript abundance was compared across all three sample types, P2 infection exhibited an intermediate phenotype: transcripts dysregulated by P18 showed concordant directionality in P2-infected tissue, but with a smaller magnitude of change (**Fig. 4B**).

**Figure 4.**
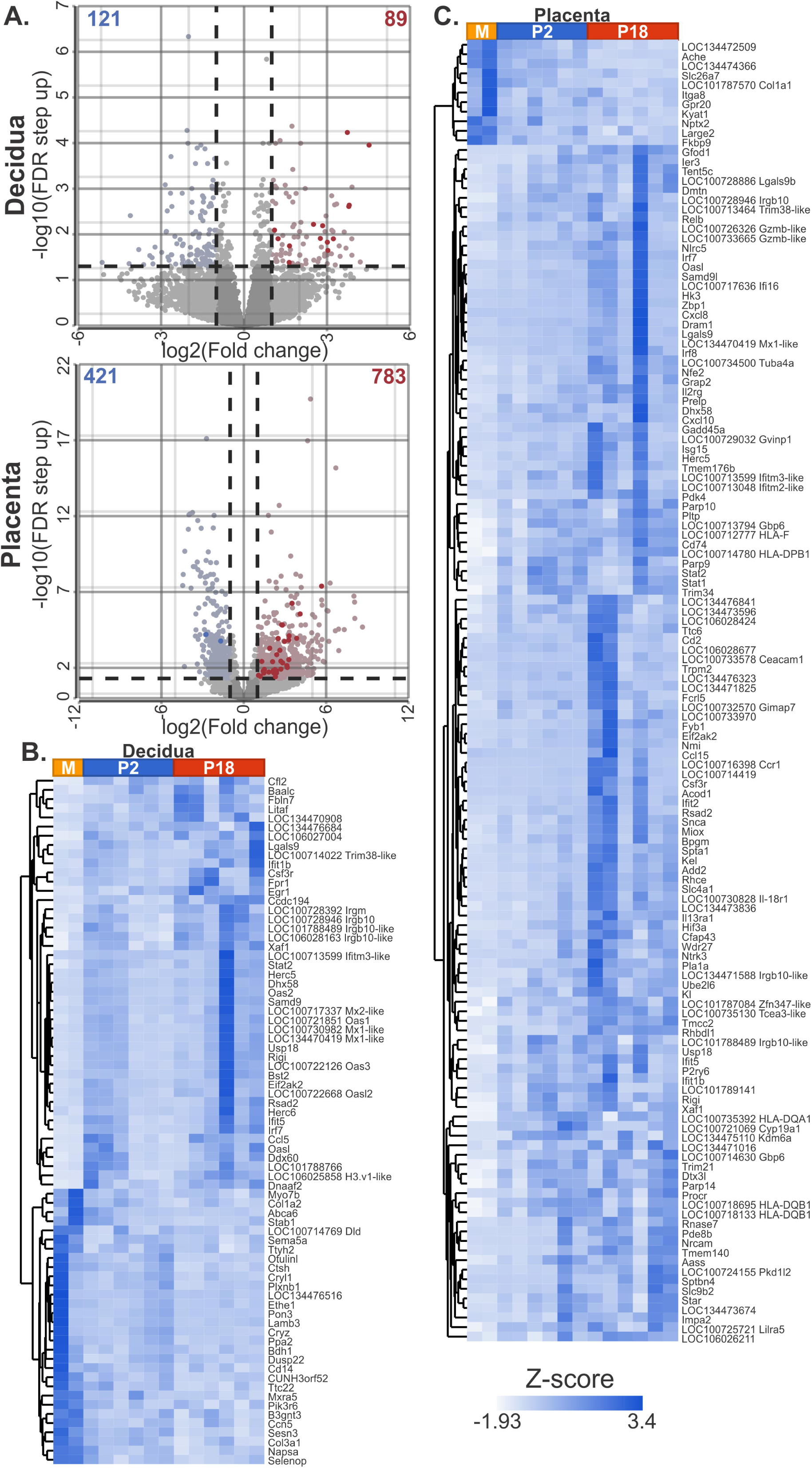
PICV infection upregulates canonical antiviral pathways in the guinea pig placenta and decidua. RNA was extracted from PICV (P2 or P18) and control (M) placenta and decidua, and transcript abundance was quantified by RNA-Seq. (**A**) Volcano plots illustrate transcripts that were significantly (≥ 2-fold, FDR ≤ 0.05) up- or down-regulated when mock- and PICV P18-infected samples were compared. Transcripts with functions related defense response to virus (GO:0051607) are emphasized, and the expression of these factors in all samples of decidua (**B**) and placenta (**C**) are illustrated by heatmap.

## Discussion

Several arenaviruses, including LASV, JUNV, and LCMV, are known to infect the placenta in clinical and experimental settings. Understanding how these zoonotic viruses affect pregnancy has been difficult due to technical barriers involved in studying highly pathogenic viruses and their infrequent clinical diagnoses. In this study, we evaluated whether PICV, an established low-biosafety model of VHF that is not pathogenic in humans, can infect placental cells and cause adverse pregnancy outcomes. We found that PICV has broad tropism for first-trimester and term human trophoblasts and that infection during guinea pig pregnancy caused fetal demise. Despite causing high viral loads in the decidua and placenta, PICV was only occasionally transmitted to the fetus. These observations led us to conclude that infection of the maternal-fetal interface plays an important role in PICV pathogenesis during pregnancy.

Maternal immunity and the innate antiviral properties of the placenta prevent most viruses from transmitting vertically during pregnancy (reviewed in [42]). Trophoblasts rely on a combination of physical defenses and the constitutive expression of antimicrobial effectors, including type III interferon, to prevent viral infection in the placenta and decidua [24]. However, the expression of pattern recognition receptors, cytokines, and interferons is tightly regulated during the course of pregnancy, and these developmentally programmed changes may affect the course of viral infection in cells and tissues at different times in gestation *in vivo* and *in vitro* [11, 43, 44].

Transcriptional profiling of PICV-infected cells and tissues revealed that the virus stimulates antiviral gene expression (**Figs. 1 and 4**). While this may seem like an expected result, there is considerable heterogeneity in how TSCs, TSC-derived trophoblasts, trophoblast organoids, and placental explants respond to diverse viruses [44-47]. PICV infection upregulates many ISGs and other factors associated with type I interferon signaling (GO:0060337) and defense responses to virus (GO:0051607). While PICV infection did not affect the transcription of interferon genes, similar patterns of ISG induction have been reported when TSCs and trophoblast organoids are infected with RNA viruses, such as Zika virus (ZIKV) or rubella virus, but not DNA viruses, like cytomegalovirus or herpes simplex virus [45-47]. In murid models, type I interferon signaling triggered by placental ZIKV infection or maternal poly I:C treatment can cause placental dysfunction and fetal wastage [22, 48, 49]. During human pregnancy, systemic lupus erythematosus is associated with increased circulating IFNα levels and high rates of pregnancy complications related to placental dysfunction [50]. Individuals with Aicardi-Goutieres syndrome and other type I interferonopathies present with a similar spectrum of CNS injuries as are observed in cases of congenital viral infection, though the effects of these syndromes on the placenta have not been examined [51]. Thus, the induction of type I interferon responses by PICV could drive placental dysfunction. Other mechanisms, such as immune cell recruitment to the infected maternal-fetal interface or disrupted hormonal signaling could also contribute to placental dysfunction. Many transcripts that play important non-immune roles in normal placental and decidual function such as extracellular matrix remodeling (*Lamb3*, *Mmp15*) and hormone signaling (*Ptgs1*,*Igfbp5*) were disrupted by PICV infection.

PICV and other arenaviruses are macrophage-trophic [19, 52]. It was recently reported that LASV could infect the reproductive tract of non-pregnant female guinea pigs, where the virus was detected in the uterus and endometrium by 4 dpi [53]. We postulate that infected macrophages seed the decidua and yolk sac with PICV at early times post infection. Viral replication in these tissue compartments may lead to a breakdown in the barrier function of the placenta, facilitating infection of the labyrinth at later time points. Differences in pregnancy outcomes caused by P2 or P18 infection may reflect P18’s capacity to cause higher systemic viral loads in both pregnant and non-pregnant animals [18]. Normal immune adaptations that occur during pregnancy can increase the severity of some viruses, such as influenza and hepatitis E, resulting in higher viral loads and more severe clinical disease [54]. However, we observed that P18 caused similar viral loads in pregnant and nonpregnant animals, leading us to conclude that pregnancy does not similarly increase the severity of PICV.

Widespread PICV infection of the yolk sac and its subsequent destruction were unexpected findings (**Fig. 3**). The fetal membranes function as an important immune barrier against ascending bacterial and viral infections, but the role of the membranes in the hematogenous transmission of viruses is less well understood [55]. Human cytomegalovirus and ZIKV can infect the chorioamnion, and guinea pig cytomegalovirus can infect the yolk sac and amnion [39, 56, 57]. In macaques, ZIKV infected cells have been detected in the chorion before virus can be observed in either the fetus or the mesenchymal tissue of placental villi [58]. Viruses could gain access to the fetal compartment by transmitting through the parietal decidua to the fetal membranes via the paraplacental route, effectively circumventing the antiviral defenses of STBs and the chorionic villi [59]. Our finding that the yolk sac is often extensively infected with PICV before the labyrinth is consistent with this model. In guinea pigs, the yolk sac involutes to form the positional equivalent of the human chorion. Where the human yolk sac functions primarily in early hematopoiesis, the guinea pig yolk sac functions as the choriovitelline placenta, a secondary organ for fetomaternal exchange [20]. Substance transport and the passive transfer of maternal antibodies by the yolk sac may be the most important for fetal development during mid gestation [60]. Destruction of the yolk sac by PICV could result in fetal anemia, growth restriction, and fetal demise.

Recent pandemics caused by ZIKV and SARS-CoV-2 illustrated the challenge of understanding how emerging viruses can affect pregnant women and their developing offspring. Given accumulating evidence that diverse arenaviruses from both the Old and New World lineages can cause adverse pregnancy outcomes, development of an experimental model for understanding host-virus interactions during pregnancy and in the placenta is needed to determine whether vaccination or antiviral therapeutics can prevent severe disease. Our findings demonstrate that PICV can be used as a low-biosafety model of VHF during pregnancy and that the virus can replicate in first-trimester and term trophoblasts. Additional studies are needed to assess whether PICV and highly pathogenic arenaviruses like LASV and JUNV can exert similar effects on the human placenta and in pregnant guinea pigs and to confirm how PICV causes placental dysfunction.

## Materials and Methods

### Key Resource Table

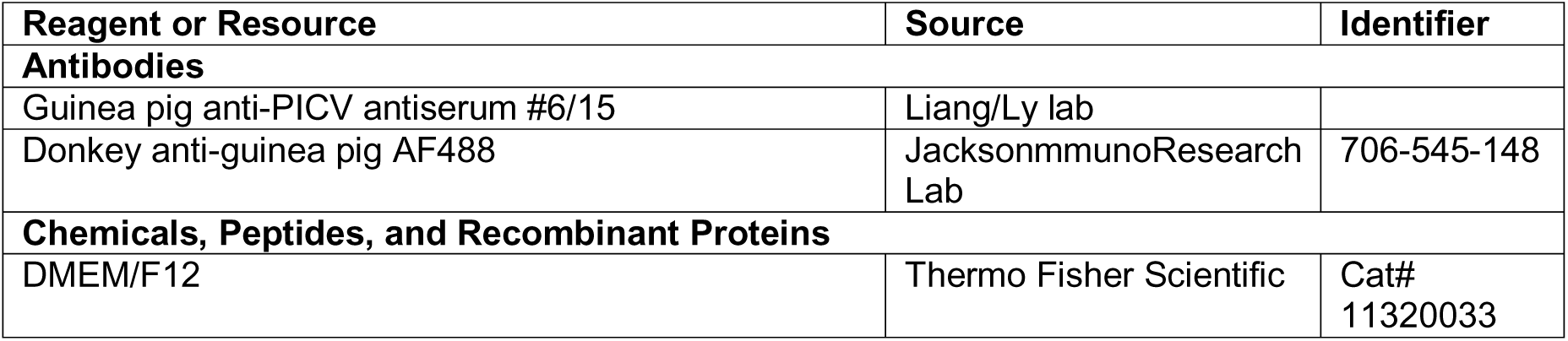
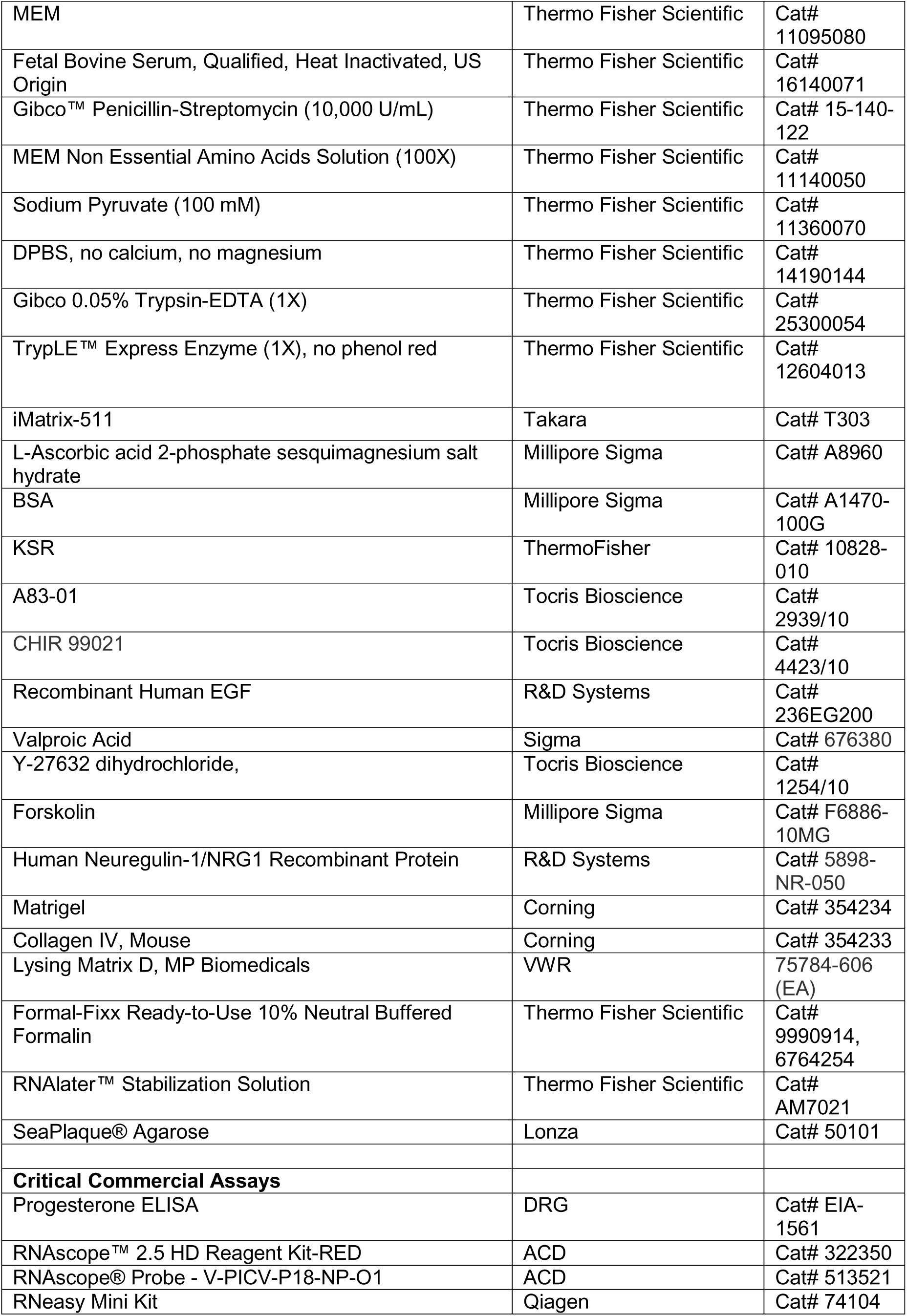
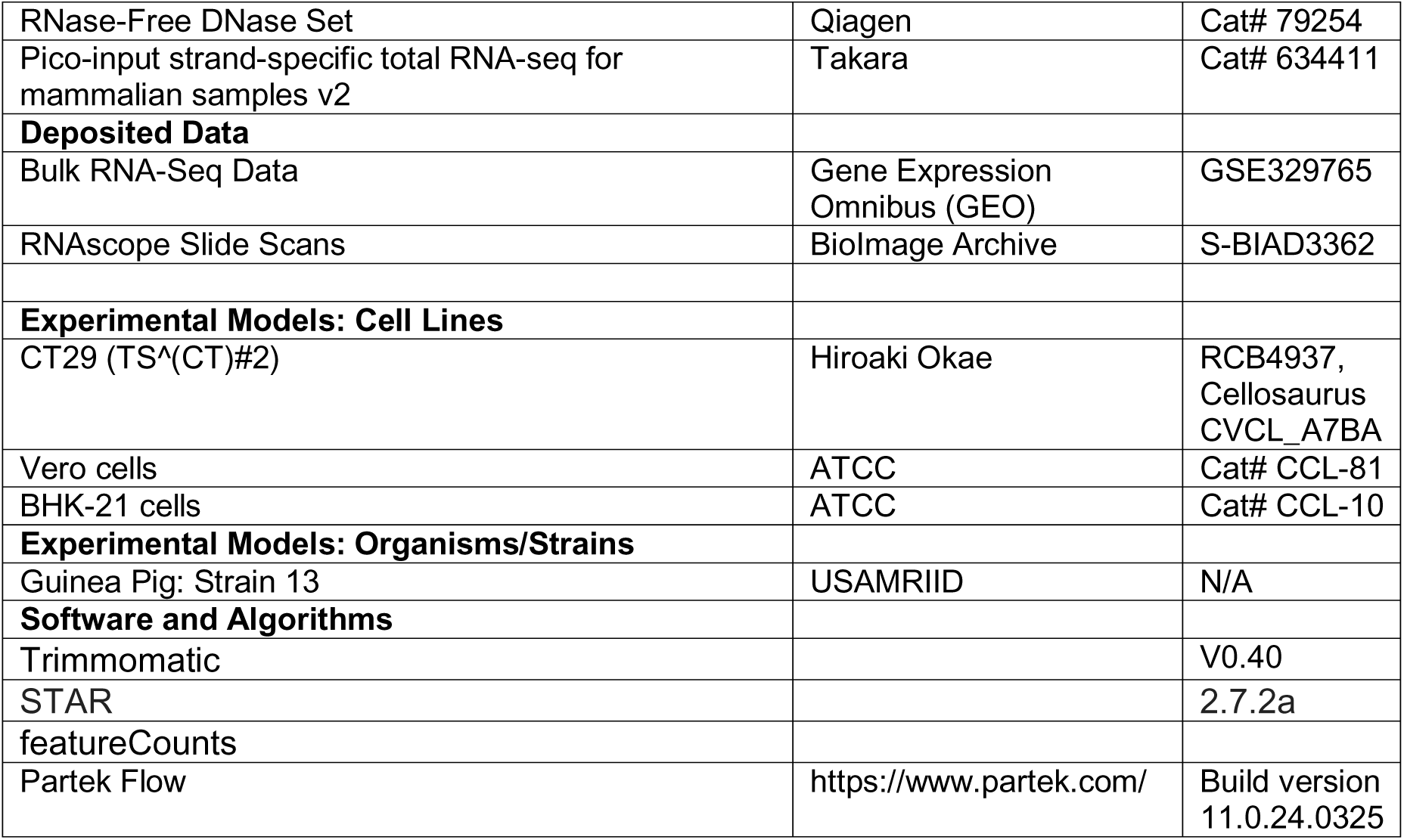

### Cells and Virus

Vero cells (ATCC CCL-81) and TSCs (CT29) were propagated as previously described [26, 38, 45]. Recombinant stocks of PICV P2 and P18 were rescued from reverse genetics systems as previously described and amplified in BHK-21 cells [61]. Plaque assays were completed on Vero cells and used to quantify infectious PICVs in viral stocks and samples collected throughout this study.

### PICV infection in human placental explants

Human placentas were obtained following informed consent by the Gestational Origins of Pediatric Health Repository (GOPHER), a tissue repository approved by the University of Minnesota Institutional Review Board (STUDY00016978 [Approved 23/01/2023], Sarah Wernimont MD PhD, Principal Investigator). Three placentas were collected from term (39-41 weeks gestation) planned repeat Cesarean births between 26/07/2023 and 28/08/2023. All participants had healthy pregnancies, no identified medical conditions, and delivered at the University of Minnesota Medical Center. Donated tissue was transported to the lab immediately following birth and processed within 45 minutes of collection. Under aseptic conditions, decidua and fetal membranes were removed, and chorionic villi were isolated and washed with PBS. Villi were dissected into ∼1 mm^3^ explants. Individual explants were placed into twelve-well plates and maintained in DMEM-F12 supplemented with 10% FBS and 1% Pen/Strep (Thermo Fisher Scientific). Explants were incubated at 37°C, 5% CO_2_ for 24 h before they were infected with 5x10^5^ PFU of P18 PICV for 1 hr. The virus-containing media were carefully aspirated and the explants were washed with DPBS three times. Samples were collected for virus quantification by plaque assay and RNAscope at 24, 48, and 72 hpi. To quantify extracellular virus, virus-containing media were centrifuged at 250x*g* to pellet debris and serially diluted. To quantify cell-associated virus in the tissue, explants were collected and homogenized in 1 ml of cell-culture media using Lysing Matrix D and a FastPrep-24 (MP Biomedicals) at 4.0 m/s for 20 s. For RNAscope, explants were fixed with FormalFix (Thermo Scientific) for 24 h and paraffin-embedded.

### PICV infection of human trophoblasts

TSCs were seeded into 12-well plates and differentiated into EVTs as previously described [45]. For STB differentiation, TSCs were seeded onto collagen IV-coated plates (Corning) and grown for 24 h with TSC complete media before the media was exchanged with STB differentiation media. EVT and STBs were infected with PICV P18 at 4 d post-seeding, TSCs were infected at 48 h post-seeding. Each well was infected with 1x10^4^ PFU of P18 (estimated MOI of 0.1). After virus had adsorbed for 1 h, the cells were washed with dPBS and incubated in cell-type specific media. Cells were scraped into the cell-culture media at 3, 24, 48, and 72 hpi and flash frozen. To measure viral loads, samples were freeze-thawed three times and plaque assays were completed as described above.

To visualize PICV infection by immunofluorescence assay, 2x10^4^ Vero cells or TSCs were seeded into each well of a µ-Slide 4 Well chamber slide (ibiTreat, ibidi). 24 h after seeding, the cells were infected with 2x10^5^ PFU of P18. After virus had adsorbed for 1 h, the cells were washed with dPBS and incubated in cell type-specific media. At 24 hpi, the cells were fixed in a freshly-prepared solution of 4% paraformaldehyde for 15 m, permeabilized with a solution of 0.2% Triton X-100 for 10 m, and blocked with 1% bovine serum albumin (BSA) in 0.1% Tween 20 for 1 h. Serum from a PICV-infected guinea pig was diluted 1:200 in 1% BSA/0.1% Tween blocking buffer and incubated with the cells for 1 h. The cells were washed three times with blocking buffer for 5 m each. Secondary donkey anti-guinea pig AF488-conjugated antibody was incubated with the cells for 1 h. The cells were washed three times with blocking buffer for 5 m. To visualize nuclei, the cells were stained with a 1:4000 dilution of DAPI (5 mg/ml stock, Thermo Fisher Scientific D1306) for 10 m. The diluted DAPI was aspirated, and the ibidi Mounting Medium was added to the wells, which were stored overnight at 4°C before imaging. All other incubations were conducted at room temperature in the dark, and images were captured using a Nikon TI-E fluorescent microscope.

### Ethics Statement

All animal procedures were conducted in accordance with protocols approved by the Institutional Animal Care and Use Committee (IACUC) at the University of Minnesota, Minneapolis (Protocol ID: 2106-39180A). Experimental protocols and endpoints were developed in strict accordance with the National Institutes of Health Office of Laboratory Animal Welfare (Animal Welfare Assurance # D16-00288), Public Health Service Policy on Humane Care and Use of Laboratory Animals, and United States Department of Agriculture Animal Welfare Act guidelines and regulations (USDA Registration # 41-R-0005) with the oversight and approval of the IACUC. Guinea pigs were housed in a facility maintained by the University of Minnesota Research Animal Resources, who are accredited through the Association for Assessment and Accreditation of Laboratory Animal Care, International (AAALAC). All procedures were conducted by trained personnel under the supervision of veterinary staff.

### PICV infection of Guinea Pigs and Tissues Collection

Two to three-month-old inbred strain 13 guinea pigs were bred [17]. Boars remained housed with dams for three days postpartum to establish second, timed pregnancies that were used for experimental PICV infections. Pregnancies were confirmed by progesterone ELISA (DRG) on or after 21 dGA and by palpation. Pregnant guinea pigs were mock- or PICV-infected at 35 dGA; age-matched virgin females were infected with PICV (N=3/group). Stocks of PICV were diluted to 2x10^5^ PFU/ml in DPBS, aliquoted, and flash frozen in single-use aliquots for animal experiments. Guinea pigs were injected subcutaneously into the scruff of the neck with 0.5 ml of diluted virus or with DPBS. Experiments were planned such that guinea pigs would be infected with similar doses of P2 or P18. Virus that remained post-infection was back titered, which revealed that P18-infected guinea pigs had received a lower infectious dose than expected (∼5000 PFU P18 and ∼60,000 PFU P2/guinea pig). Guinea pigs were euthanized immediately after delivering pups or at 10, 15, or 20 dpi. Placenta and decidua were collected, divided, and portions were formalin-fixed and embedded in paraffin, stabilized in RNAlater (Thermo Fisher Scientific), or frozen and stored at -80°C. Other maternal and fetal tissues were stabilized in RNAlater or frozen and stored at -80°C. To quantify viral load in guinea pig tissue samples, samples were homogenized in 1X dPBS using a Bead Bug homogenizer at 400x10 for 40 seconds.

### *In situ* hybridization

5-μm sections of formalin-fixed, paraffin-embedded tissue were mounted onto Superfrost Plus slides (ThermoFisher). After air drying the tissue sections overnight, the slides were baked at 60 °C for 1 h. Tissue was deparaffinized and pretreated using the recommended standard protocol for RNAscope^®^ 2.5 Assays (ACD Document #322452). For target retrieval, samples were incubated at 99°C for 15 minutes, and the slides were treated with RNAscope Protease Plus for 30 m. Slides were stained using the RNAscope 2.5 HD Detection Reagent – RED (ACD Document # 322360-USM) the V-PICV-P18-NP-O1 RNAscope probe, which targets NP mRNA and antigenomes. Slides were scanned using a Huron TissueScope LE. Image data are available in the BioImage Archive (http://www.ebi.ac.uk/bioimage-archive) under accession number S-BIAD3362 [62].

### RNA Sequencing

For RNA sequencing, TSCs were seeded at a density of 1.6X10^5^ cells/well to 6-well plates 24 h before mock- or P18-infection (5X10^6^ PFU/well, approximate MOI of 10). RNA was extracted from two replicates for each condition at 24 hpi using a RNeasy Mini Kit (Qiagen) with an on-column DNase digest according to the manufacturer’s instructions. RNA was extracted from guinea pig decidua and placenta (10 dpi, 2 samples from each litter plus two samples from a gestational age-matched uninfected control) using a RNeasy Mini Kit (Qiagen). About 30 mg of RNAlater-stabilized tissue was homogenized in 0.6 ml of β-mercaptoethanol-containing RLT Buffer using lysing matrix D tubes (MP Biomedicals) and a FastPrep 24 bead beater (MP Biomedicals) by pulsing the samples once at 6 m/s for 30 s. The homogenized samples were centrifuged at 28,000x*g.* The cell lysate was transferred into a new microcentrifuge tube, and 450 μL of lysate was mixed with 1X volume of 70% ethanol. The standard RNeasy protocol with on-column DNase digest was used to purify RNA after this step. RNA quality was assessed using an Agilent 4200 TapeStation. While the RNA extracted from PICV-infected TSCs was of high quality (RIN=10.0), the quality of RNA extracted from decidua and placenta was more variable (RIN scores ranging from 5.1 to 9.1). RNA sequencing libraries were prepared using the Pico-input strand-specific total RNA-seq for mammalian samples v2 (Takara, mean quality scores for all libraries ≥Q30.). Libraries were sequenced using an Illumina NovaSeq (2x150-bp run format, >4X10^7^ reads/sample).

Sequencing reads were processed using Trimmomatic set to default parameters [63] and aligned to reference files that contained either the human (GRCh38) or guinea pig (mCavPor4.1, GCF_034190915.1) genome with STAR version 2.7.2a using the default parameters [64, 65]. Gene count tables were prepared with featureCounts and imported into Partek Flow (version 11.0.24.0325) for subsequent analyses [66]. Features with a maximum count ≤ 1.0 were excluded, transcript abundance was normalized using the median ratio method, and DESeq2 was used to identify differentially regulated transcripts. Mock- and PICV-infected TSCs were compared, as were mock- and PICV-infected samples of guinea pig placenta and decidua. Differentially regulated features were defined as having a ≥2-fold change in abundance and an FDR ≤ 0.05. The abundance of select transcripts was visualized by volcano plot and heatmap, and gene set enrichment (completed either with Partek Flow’s algorithm [human samples] or g:Profiler [guinea pig samples]) was used to identify processes that were affected by infection [67, 68]. Sequencing data has been uploaded to the Gene Expression Omnibus (GEO) (ascension number GSE329765).

## Acknowledgments

This project was funded by a University of Minnesota College of Veterinary Medicine 2023 Intramural Grant (HL and CJB) and the Eunice Kennedy Shriver National Institute of Child Health and Human Development (R01HD109252 to CJB and R01HD117756 to CJB, HL, and YL). Additional support was provided by the National Institutes of Health’s National Center for Advancing Translational Sciences (UM1TR004405), which provided funding for the Biorepository and Laboratory Services Team and funding for JSH (5T32TR004385), and the resources and staff at the University of Minnesota University Imaging Centers (SCR_020997) and the Genomics Center (https://genomics.umn.edu). The content of this article is solely the responsibility of the authors and does not necessarily represent the official views of the National Institutes of Health.

## Author Contributions

Conceptualization: CJB, HL, and YL. Methodology: CJB, HL, and YL. Investigation: CJB, HM, JSH, QH, TBR, BF, PC, SAW, and TKM. Data Curation: CB and MG. Funding Acquisition: CJB, YL, and HL. Writing—Original Draft: CJB. Writing—Reviewing and editing: CJB, YL, HL, HM, JSH, QH, TBR, PC, SAW, and TKM. Supervision: CJB, HL, and YL.

## Declaration of Interests

The authors declare no competing interests.

## Supporting information

**S1 Table. Viral load data and pathologic findings from guinea pig infections.**

**S1 Figure. Change in guinea pig weight after PICV infection.** Time-mated and non-pregnant (NP) guinea pigs were infected with PICV. The change in each animals’ weight observed after infection with (**A**) P18 or (**B**) P2 was plotted. **×** indicates the day when P18-infected dams delivered stillborn pups.

## Notes

https://www.ncbi.nlm.nih.gov/geo/query/acc.cgi?acc=GSE329765

https://www.ebi.ac.uk/biostudies/bioimages/studies/S-BIAD3362

### Competing Interest Statement

The authors have declared no competing interest.

### Summary of Updates

Supporting information has been uploaded with the manuscript. Funding information and supporting datasets have been associated with the preprint.

