## Supplementary figures and images for "Pichinde virus models intrauterine infection by hemorrhagic fever-causing arenaviruses"

### Figure S1

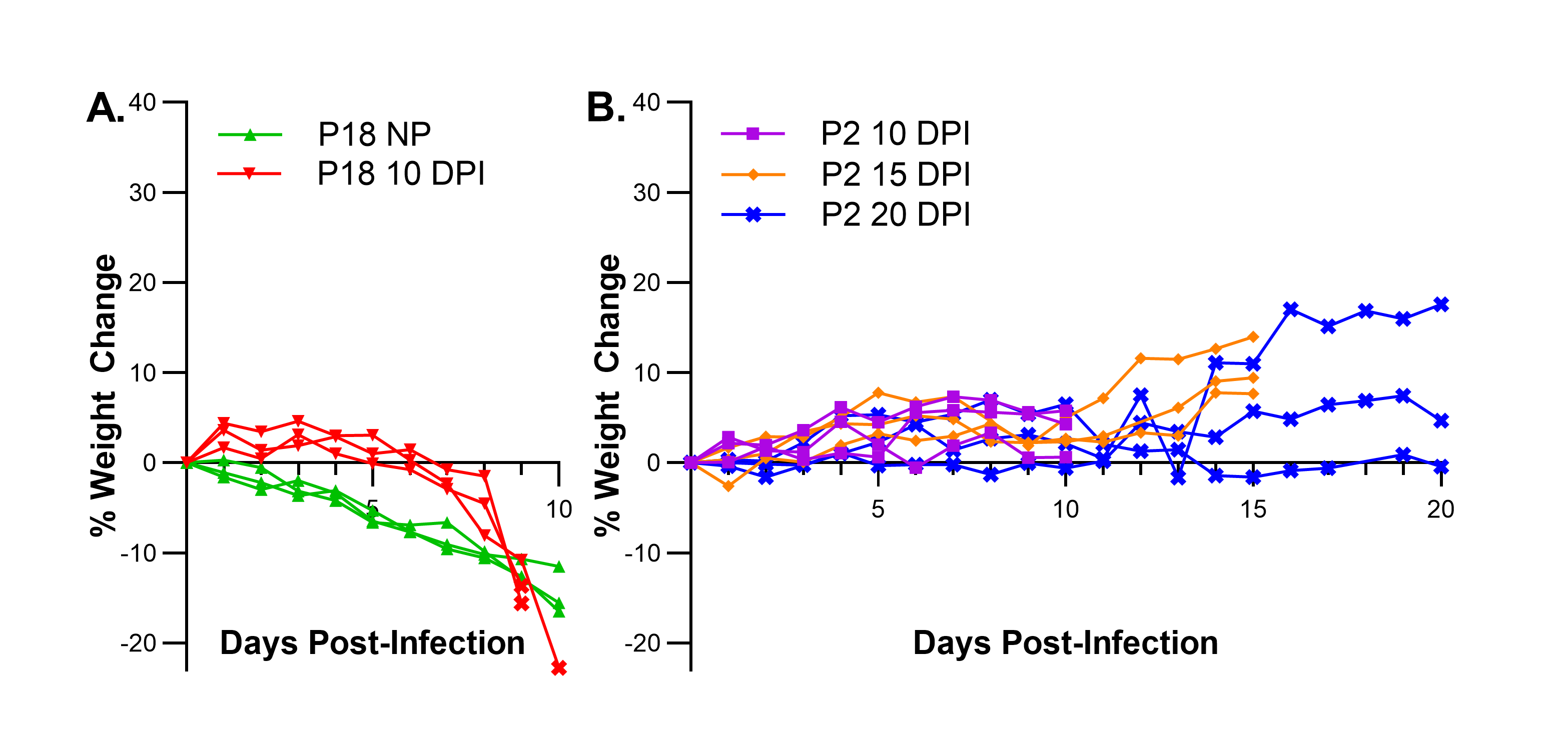
